# Adjuvant effects of multifunctional transcription factor and BCG target YB-1: exogenous YB-1 enhances specific antibody production *in vivo* and protects mice against lethal *E. coli* challenge

**DOI:** 10.1101/2022.11.09.515841

**Authors:** Anna O. Shepelyakovskaya, Ludmila Alekseeva, Elena A. Meshcheryakova, Khanafiy Boziev, Alexandra Tsitrina, Vadim T. Ivanov, Fedor Brovko, Yuri Kotelevtsev, Richard Lathe, Alexander G. Laman

**Affiliations:** Shemyakin and Ovchinnikov Institute of Bioorganic Chemistry, Pushchino Branch, Russian Academy of Sciences, Pushchino, Moscow Region 142290, Russia; Shemyakin and Ovchinnikov Institute of Bioorganic Chemistry, Russian Academy of Sciences, Moscow 117997, Russia; Optical Research Group, Koltzov Institute of Developmental Biology, Russian Academy of Sciences, Moscow 119991, Russia; Skolkovo Institute of Science and Technology, Moscow 121205, Russia; Division of Infection and Pathway Medicine, University of Edinburgh, Edinburgh EH16 4SB, UK

**Keywords:** adjuvant, BCG, GMDP, muramyl peptide, sepsis, vaccine, YB-1, YBX1

## Abstract

There is growing interest in the beneficial effects of immune system boosting through the administration of adjuvants, not only in acute infections such as COVID but also in chronic degenerative disorders that are potentially associated with infection. The best-known immunopotentiators are Freund’s complete adjuvant (FCA) and its relative Bacille Calmete–Guérin (BCG), both based on *Mycobacterium* species. The key pathogen-associated molecular patterns (PAMPs) in both FCA and BCG are muramyl dipeptides (MDPs and glucosaminyl-MDP, GMDP). We previously identified the evolutionarily conserved protein Y-box factor YB-1/YBX1 as a primary target for MDP/GMDP. Unlike other host receptors for PAMPs, YB-1 is a diffusible molecule, and we therefore explored whether *in vivo* administration of YB-1, rather than its PAMP ligands, might enhance the immune response to a bacterial antigen and/or influence survival in the face of bacterial infection. We report that mice receiving YB-1 plus GMDP *in vivo* mount a significantly increased B cell response versus GMDP alone against a test antigen (*Yersinia pestis* V antigen), and that YB-1 administration alone significantly promotes survival in the face of lethal bacterial (*Escherichia coli*) challenge *in vivo*. Independent confirmation is warranted because recombinant YB-1 and its ligands could hold great promise both as adjuvants and as therapeutics.

## INTRODUCTION

Vaccine efficacy is markedly enhanced by stimulatory molecules such as aluminium salts (alum), lipopolysaccharide (LPS), and muramyl dipeptides (MDPs). These adjuvants are thought to bind to cellular receptors that, in turn, promote the expression of cytokines and other molecules that further enhance antigen-specific immune responses. However, only four adjuvant molecules are currently licensed for human use. The most common is aluminium hydroxide and related aluminium salts, known as ‘alum’; more recently licensed adjuvants include lipid-based molecules (monophosphoryl lipid A, MF59/squalene), a triterpene glycoside (QS-21), and CpG 1018 (a synthetic bacterial DNA mimetic) (https://www.cdc.gov/vaccinesafety/concerns/adjuvants.html; [1]). However, alum remains the key adjuvant molecule in >90% of vaccines, and there is an urgent need to develop new immunopotentiating agents [2].

The most powerful adjuvant known, Freund’s complete adjuvant (FCA), is based on heat-killed *Mycobacterium tuberculosis* in mineral oil [3]. Nevertheless, the potency of FCA in inducing a local immune reaction is so high that it can induce lesions at the site of inoculation, and is therefore not approved for human use [3]. By contrast, Bacille Calmete–Guérin (BCG), an anti-tuberculosis vaccine based on attenuated *Mycobacterium bovis*, is approved for human use and is highly effective against tuberculosis [4]; importantly, BCG shares many of the non-specific immunostimulatory effects of FCA, and BCG inoculation has been observed to confer protection against unrelated infectious agents including viruses [5,6], malaria [7], and respiratory infections [8–10].

Two of the most powerful adjuvants, FCA and BCG, are therefore both based on *Mycobacterium species*. Because of the toxicity of FCA, early investigators sought to determine the key molecules (pathogen-associated molecular patterns, PAMPs) involved, leading to the identification of muramyl dipeptide (MDP; *N*-acetyl muramic acid attached to a short L-Ala-D-isoGln amino acid chain) and its close relative, acetylglucosaminyl-MDP (GMDP; that contains an additional carbohydrate group), as the active bacterial cell-wall peptidoglycan component(s) that can replace mycobacterial cells in FCA ([11–13], reviewed in [14]). Because GMDP is considered to be less toxic than MDP [15], many studies have focused on this molecule. The adjuvant effects of muramyl peptides in protecting against challenge infection with different bacteria (e.g., [16,17]) and viruses (e.g., [18–20]) have been confirmed (reviewed in [21]).

Two host molecules have been identified as receptor targets for muramyl peptide PAMPs – NOD2 and YB-1 [22–25]. Whereas NOD2 and its relative NOD1 (that responds to D-glutamyl-*meso*-diaminopimelic acid [26]) are intracellular receptors, YB-1 differs in that it is a diffusible multifunctional transcription factor [27,28] that is present not only in the nucleus but also in the cytosolic and extracellular compartments [29]. Notably, blood levels of YB-1 are remarkably elevated in patients with sepsis [30], suggesting that YB-1 upregulation may be a protective response to infection, and raising the question of whether YB-1 itself, with or without GMDP/MDP ligand, might have immunostimulatory/adjuvant action.

Biologically active YB-1 can be produced in pharmacological amounts in recombinant bacteria [24,25,31]. In this paper we investigated whether administration of recombinant YB-1 *in vivo*, with or without GMDP, might enhance the immune response to a specific antigen and/or influence survival in the face of acute infectious challenge.

## METHODS

### Reagents

GMDP (*N*-acetylglucosaminyl-*N*-acetylmuramyl-alanyl-D-isoglutamine) was synthesized by Peptek (Moscow, Russia). Recombinant YB-1 (324 amino acids, rabbit, that differs at only three positions from human YB-1, 99% homology; and at eight positions from mouse YB-1, 98% homology) that was expressed in *E. coli* and purified as described previously [31], was kindly provided by Professor L.P. Ovchinnikov. *Yersinia pestis* V antigen (also expressed in *E. coli*) was kindly provided by Professor A.P. Anisimov at the State Research Center for Applied Biotechnology and Microbiology (Obolensk, Russian Federation).

### Antibody response boosting by YB-1 and GMDP

Five groups of mice (BALB/c mice, *N* = 3 in each case) were immunized with (i) control PBS only, (ii) *Y. pestis* V antigen (5 μg/mouse), (iii) V antigen (5 μg/mouse) + YB-1 (1 μg/mouse), (iv) V antigen (5 μg/mouse) + GMDP (10 μg/mouse), (iv) V antigen (5 μg/mouse) + YB-1 (1 μg/mouse) + GMDP (10 μg/mouse). Spleens were collected after 2 weeks, mononuclear spleen cells were purified (Ficoll density-gradient centrifugation), and cultured *in vitro* (DMEM plus 10% FBS) for a further 48 h in the presence or absence of the same additives or combinations of additives as before (V antigen, 5 μg/ml; YB-1, 1 μg/ml; GMDP, 10 μg/ml). Cells were harvested, fractions from each of the 5 groups were pooled, counted, and 10^5^ cells from each pool were subjected to analysis as follows. To prevent subsequent antibodies interacting with Fc receptors, 1 μg of purified rat anti-mouse CD16/CD32 was added to each pool (Mouse BD Fc Block, BD Biosciences, cat. 55142) and incubated for 30 minutes at 4°C. B cells were identified following incubation with antibodies against the pan B cell marker CD45R {peridinin-chlorophyll protein (PerCP)-labeled rat anti-mouse CD45R/B220, BD Pharmingen, cat. 553093} and against IgM/IgD {PE-labeled rat anti-mouse IgM, BD Pharmingen, cat. 553517; phycoerythrin (PE)-labeled rat anti-mouse IgD, BD Pharmingen, cat. 558597}. To determine specific V antigen responses, cells were incubated with FITC–V antigen (1 h in the dark at 4°C) and the antigen-specific B cell population was identified by flow cytometry using a BD ACCURI C6 flow cytometer (BD Biosciences). The non-parametric Kruskal–Wallis test was used to compare the medians (*H* statistic) of the responses.

### *In vivo* acute toxic challenge

BALB/c mice were maintained at the Institute of Bioorganic Chemistry, Pushchino, and housed under conventional conditions. Groups of 8-week-old male mice were injected intraperitoneally (i.p.) with *E. coli* TG-1 in PBS at a dose 5 × 10^7^ bacteria per animal that in preliminary experiments caused the death of >90% of animals. Two experimental groups (each *N* = 18) were treated with i.p. 1 μg YB-1/animal in PBS 12 h before or concomitantly with challenge; control animals received PBS instead of *E. coli*. Mortality assessment included both frank demise and overt morbidity sufficient to justify euthanasia on welfare grounds. Statistical analysis of Kaplan–Meier survival curves employed both *t*-testing and log-rank testing (Statistics Kingdom: https://www.statskingdom.com/kaplan-meier.html).

### Animal procedures

Animals were obtained from the Laboratory Animal Breeding Nursery, Pushchino Branch, Institute of Bioorganic Chemistry, Russian Academy of Sciences, which is accredited by the AAALACi (Association for Assessment and Accreditation of Laboratory Animal Care International). All regulated procedures were conducted in accordance with Russian Academy of Science Guidelines for Animal Experimentation; the specific protocols and ethical permissions for this work were authorized under registration number 852/21.

## RESULTS

### Synergistic adjuvant activity of YB-1 and GMDP

To address whether administration of YB-1 might stimulate B cell immunity, we immunized mice with a control bacterial antigen (*Yersinia pestis* V antigen; 5 μg/mouse) in the presence or absence of recombinant YB-1 (1 μg/mouse) +/− GMDP (10 μg/mouse). Two weeks later mononuclear spleen cells were isolated, cultured (48 h, with the same agent), and the specific B cell population was analyzed by flow cytometry (Figure 1A). Labeling with fluorescent (FITC)-labeled V-antigen was used to determine the specific antigen-binding activity of surface immunoglobulin.

**Figure 1.**
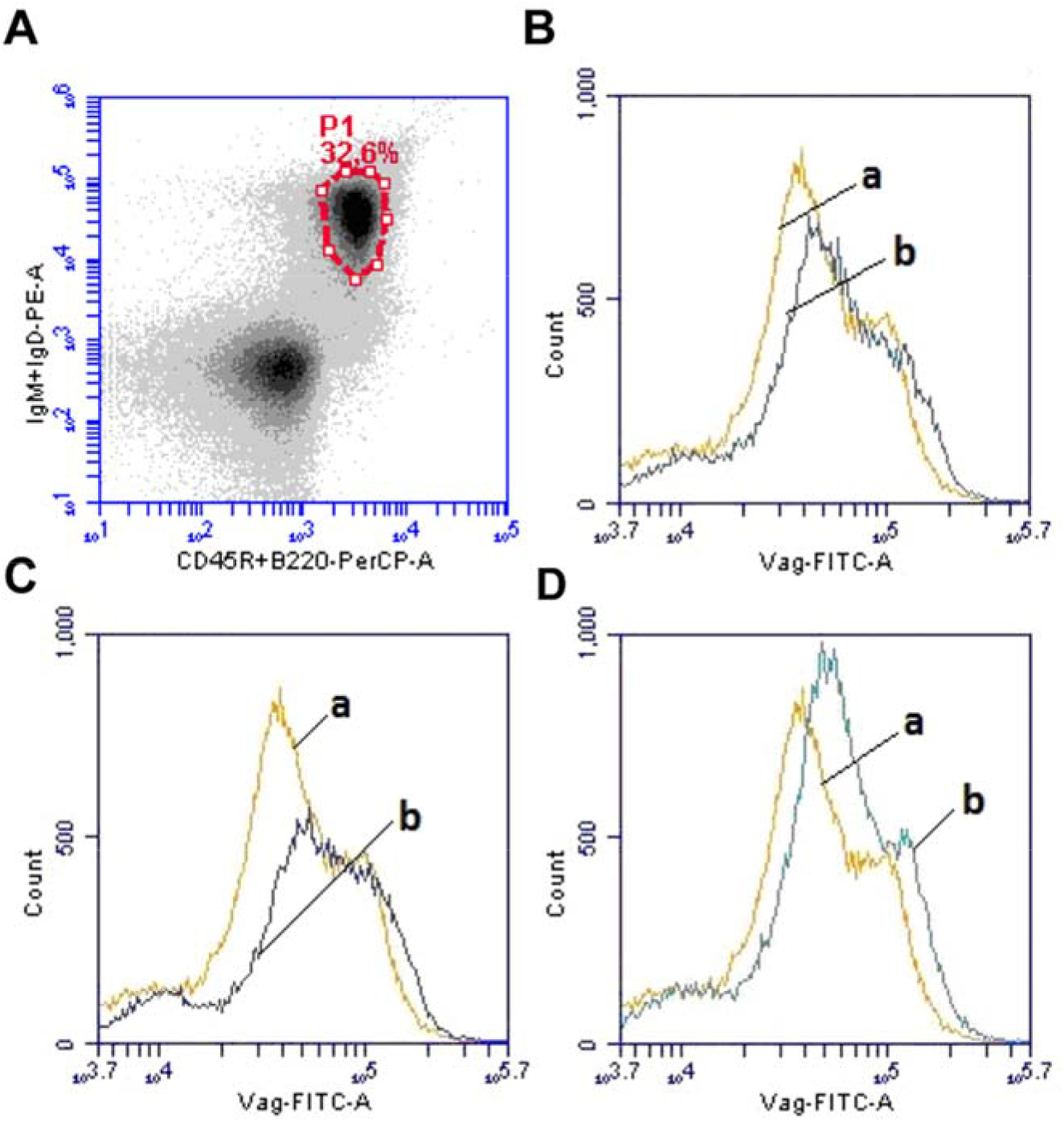
Adjuvant effects of recombinant YB-1 and GMDP. (A) Characterization of mouse B cell responses by cell sorting. Mice (*N* = 3 in each case) were immunized with control bacterial antigen (*Yersinia pestis* V antigen) in the presence or absence of YB-1 and GMDP, mononuclear spleen cells were collected after 2 weeks, boosted in culture (48 h), pooled, and analyzed by flow cytometry with antibodies against the pan B cell marker CD45R and surface immunoglobulins (IgM/IgD). The panel shows the proportion of cells (fraction P1) stained with antibodies against B cell marker CD45R/B220. (B–D) Antigen reactivity (cellular fluorescence with bound FITC–V antigen, Vag-FITC-A) of B cell fractions from mice (*N* = 3 in each case) receiving V antigen in the presence or absence of (B) GMDP, (C) YB-1, and (D) GMDP plus YB-1. The yellow trace (a) in each case is from mice immunized only with V antigen. Note that the apparent bimodality is a common artefact of flow cytology [32]; plotted on a linear scale only one peak is present. Unimmunized control mice (PBS) showed low-level reactivity where no intensity value exceeded 10^3.5^ (not depicted). Blue curves (b) in each case are from mice immunized in the presence of GMDP (B), YB-1 (C), or both (D), showing that GMDP improves the antigen-specific response, and the synergistic effects of YB-1 plus GMDP. The antibody response with YB-1 plus GMDP coadministration (D) was significantly (*P* = < 0.001) higher than with GMDP alone (B).

Administration of GMDP together with V antigen led to a significant increase in B cell reactivity to FITC–V antigen (Figure 1B–D). Coadministration of YB-1 with GMDP produced the biggest enhancement of antigen-specific B cell reactivity in terms both of B cell number and per-cell antigen binding, indicating that YB-1 and GMDP act synergistically (Figure 1). Because the data are not normally distributed, the non-parametric Kruskal–Wallis test was used to compare the responses (*N* = 3 in each case) in Figure 1B–D. It is important to note that the apparent bimodal distribution of values into two peaks is a common artefact of flow cytometric analysis [32]. In this analysis, V antigen responses upon supplementation with YB-1 and/or GMDP (blue curves in Figure 1) were significantly different (*P* < 0.03) from V antigen alone (yellow curves in Figure 1) for B (GMDP) and D (GMDP + YB-1), whereas the increased response in C (for YB-1 in the absence of GMDP) fell short of statistical significance. Importantly, the V antigen response to the combination of YB-1 plus GMDP (blue curve in Figure 1D) was very significantly different (*P* = < 0.001) from GMDP alone (blue curve in Figure 1B).

These findings demonstrate that the bacterial cell-wall derivative GMDP potentiates the B cell response to *Y. pestis* V antigen. Although administration of YB-1 alone, a major binding target for muramyl peptides, had only a marginal effect on the antibody response, coadministration of YB-1 with GMDP markedly increased B cell number/affinity for V antigen versus GMDP alone. GMDP therefore appears to be essential for YB-1 adjuvant action, and the adjuvant effects of exogenous GMDP may therefore be ascribed to targeting of endogenous receptor proteins. By contrast, the immune response was substantially improved by adminstration of exogenous YB-1 together with GMDP, suggesting that endogenous levels of YB-1 may be suboptimal.

### Recombinant YB-1 protein protects mice from lethal *E. coli* challenge

To address whether YB-1 itself can exert adjuvant action *in vivo*, we examined whether YB-1 alone might modulate disease course in mice lethally challenged with *E. coli*, a bacterium likely to release MDP/GMDP or related peptidoglycan molecules during the course of infection. Mice were inoculated with a dose of *E. coli* titrated to give >90% lethality within 5 days post-challenge. Animals either received recombinant YB-1 (1 μg YB-1/animal) at time of challenge or were pretreated with the same dose of YB-1 12 h before challenge.

Administration of YB-1 at time of challenge had no effect on survival, and all mice succumbed to sepsis within 3–4 days. By contrast, preadministration of YB-1 12 h before challenge conferred significant protection against *E. coli* challenge, and 9 of 18 animals (50%) survived to 30 days post-challenge (*P* = <0.0001 by t-test; *P* = <0.008 and *P* = <0.001 by log-rank testing) (Figure 2).

**Figure 2.**
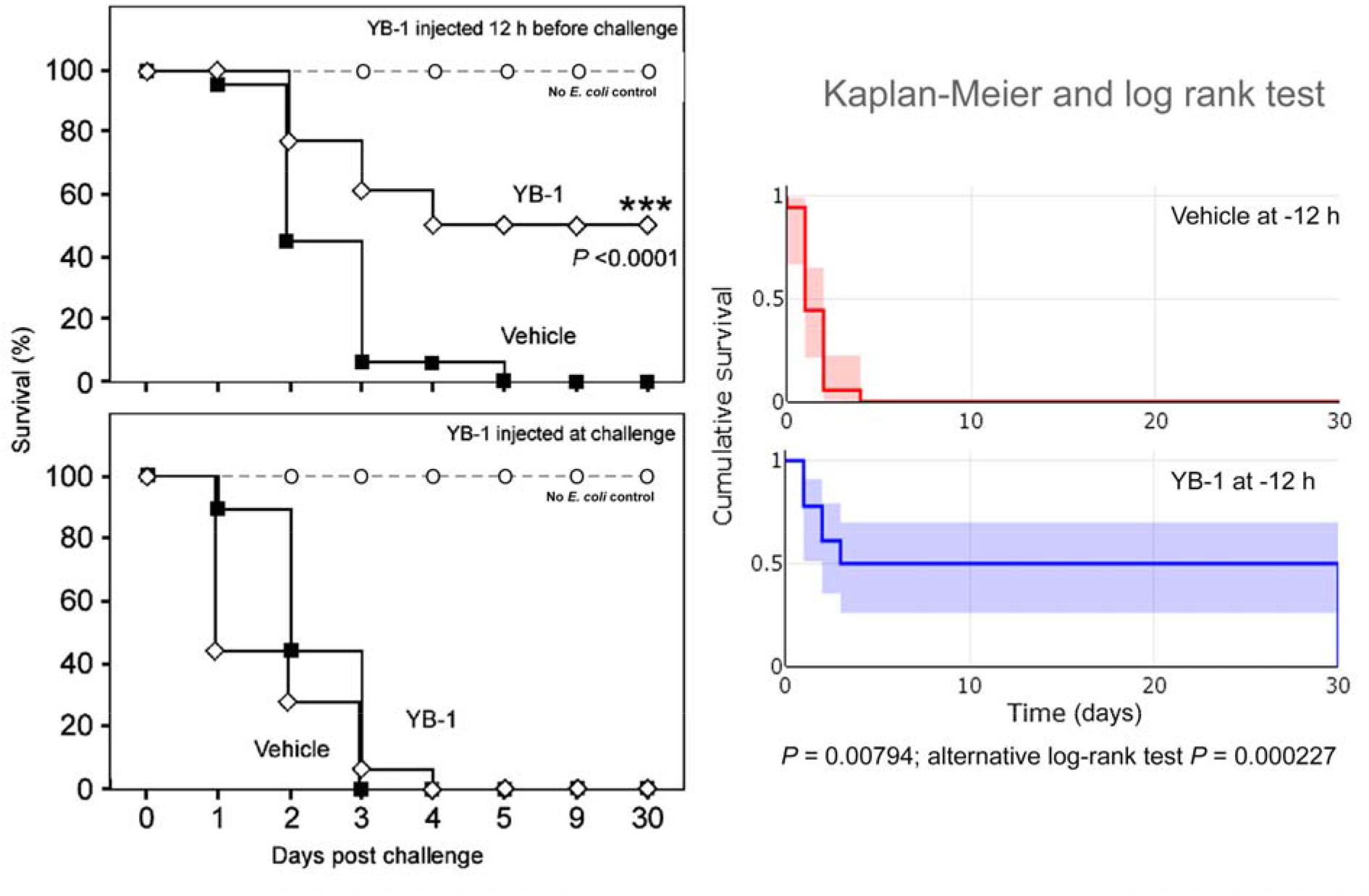
Host factor YB-1 alone can protect against lethal *Escherichia coli* challenge. Control mice were treated with YB-1 or vehicle but received PBS instead of *E. coli* challenge. (Left) Percentage survival rates of mouse groups (each *N* = 18) receiving YB-1 either 12 h before (upper panel) or concomitantly with (lower panel) lethal challenge with *E. coli* (data enlarged to show the response over days 0–4). The survival increase with YB-1 was significant versus vehicle (*P* = <0.0001 by *t*-test). (Right) Survival data from the left-hand panel replotted according to Kaplan–Meier and analyzed using log-rank testing; the survival difference was also significant in the both the standard and alternative tests (*P* = <0.008 and *P* = <0.001). Colored areas are 95% confidence intervals.

## DISCUSSION

### Augmented antibody response against *Yersinia pestis* V antigen

Our observation that YB-1 + GMDP synergistically increase the antibody response to *Y. pestis* V antigen versus GMDP alone or vehicle is an important finding. V antigen is a representative of a highly homologous family of proteins in Gram-negative bacteria that constitute the main structural element of the type III secretory system (T3SS) [33]. Although antibodies to V antigen are able to protect animals in a model of bubonic plague [34], infection with *Y. pestis* and related organisms is characterized by the development of chronic disease with a low level of immune response [35]. The search for innate immunity regulators capable of activating the response to V antigen is therefore of considerable scientific and medical importance.

### Protection against lethal *E. coli* challenge

YB-1 alone was able to promote survival in mice receiving lethal challenge injection with *E. coli*, a model of acute bacterial sepsis, and recombinant protein administration to mice 12 h before lethal challenge with *E. coli* led to 50% long-term survival. Although 12 h pretreatment (i.p.) was necessary to confer protection against lethal *E. coli* challenge, this timecourse of action is more rapid than was previously encountered. In earlier experiments a minimum period of 14 days of pretreatment with non-lethal GMDP was necessary to confer protection against lethal (LPS) challenge; protection with the GMDP analog GMPS required a minimum of 6 days pretreatment [17]. We report significant protection by administration only 12 h before lethal challenge, a highly encouraging result suggesting that further improvement might be achieved by adjustment of the dosage and/or route of administration.

### Immune boosting, cross-protection, and the wider potential of YB-1 and GMDP

The development of new adjuvants to boost vaccine efficacy, as well as to improve therapeutic treatments for both acute and chronic disease, is of outstanding current interest. In this context there has been growing interest in the use of vaccines and/or vaccine components as immunostimulatory adjuvants in protecting against unrelated infectious diseases. Forster and Abshier reported in 1937 that individuals receiving smallpox vaccine (vaccinia virus) often displayed regression of lesions caused by unrelated herpes viruses [36], a finding confirmed in some studies [37,38], although queried in others [39]. Therapeutic effects against herpes have also been reported with attenuated poliovirus-based vaccine [40,41]. The non-specific effects of vaccines in protecting against unrelated infectious agents have been extensively reviewed [42–45].

More recently it has been speculated that non-specific boosting of the immune system with BCG might protect against COVID-19 ([46] for review). This is supported by a series of recent reports. In mice, intravenous administration of BCG protected mice against lethal challenge infection with SARS-CoV-2 [47]. Medical staff reimmunized with BCG showed a significantly (*P* = 0.004) lower rate of COVID-19 infection [48]. Recent meta-analysis confirms that the risk of COVID-19 infection is significantly lower (0.61) in BCG-vaccinated individuals [49], and a consistent association was found between BCG vaccination and reduced COVID-19 severity across 21 European and North American countries (*P* = < 0.00001) [50].

Immune boosting may also have protective effects in Alzheimer’s disease (AD), a disorder that has increasingly been associated with infection [51]. Previous exposure to a range of vaccines (diphtheria, tetanus, poliomyelitis, influenza) was reported in 2001 to be associated with a 40% reduced risk of AD development [52]. More recently, in a Phase II trial of an AD vaccine, Affiris AG reported that administration of the adjuvant, alum, to control patients retarded AD-related cognitive decline across multiple measures [53,54]. Importantly, in a study of patients with bladder cancer, where BCG administration is the standard of care [55], the risk of developing AD was reported to be 4.8-fold lower in those receiving BCG than in untreated patients [56]. Reduced AD risk has also been reported for recipients of other vaccines including herpes zoster [57–59] and influenza [60–62]. In a screen of 744 routinely administered medications, four were protective against AD, and all were vaccines [63]. BCG has also been reported to attenuate adverse events in another neurological disorder, multiple sclerosis [64]. In all these cases the mechanism of protection is presumed to be via non-specific activation of the immune system through adjuvant action. In this context, YB-1 and/or GMDP/MDP may represent a new weapon in our armory that could potentially attenuate neurological disease.

In sum, our findings demonstrate that exogenous administration of a host immunomodulatory transcription factor, YB-1, can boost *in vivo* B cell responses; moreover, short (12 h) pre-exposure to YB-1 was found to protect against acute lethal challenge with *E. coli*. Because YB-1 is a host protein unrelated to the challenge organism (*E. coli*), this indicates that its effects are achieved through immunomodulation. Although YB-1 administration has been explored in other experimental models (e.g., [65]), to our knowledge this is the first demonstration that administration of a PAMP receptor (rather than its ligand) can itself enhance the immune response to antigen inoculation and protect against infectious challenge. The immunomodulatory mechanisms of YB-1 protection against lethal infectious challenge are the subject of ongoing investigations.

Further experiments will be necessary to determine whether administration of YB-1, independently of or in conjunction with its ligands GMDP/MDP, can influence cytokine secretion and offer significant protection not only against acute infections of outstanding topical interest (i.e., COVID [66]) but also against protracted infections such as those implicated in neurodegenerative diseases including AD and multiple sclerosis.

Our experiments employed 1 microgram of YB-1 per mouse, which would correspond to ~2.5 mg in human. This is to be compared to clinical dosages of therapeutic humanized antibodies that are typically in the range 5–150 mg. There is thus considerable scope to investigate higher doses of YB-1 for potential clinical use. In addition, YB-1, at 324 amino acids (aa), is bigger than commonly administered proteins such as insulin (21 aa; precursor 100 aa), interferon gamma (126 aa), and growth hormone (191 aa). Nevertheless, coagulation factor IX (Christmas factor), that is routinely administered for hemophilia B, is substantially larger (415 aa), and antibody heavy chains are in the range of 450 aa. Native human YB-1 may thus be of direct clinical utility, although the identification of shorter forms that preserve the immunopotentiating activity of the protein might expedite clinical acceptance in the management of acute and chronic infections.

In addition, the acute *E. coli* infection model employed here is to some extent a model of sepsis, a challenging condition for which there is no effective treatment beyond supportive management [67,68]. YB-1 and its ligands warrant investigation in classical models of sepsis such as cecal ligation and puncture (CLP) [69], as well as in clinical sepsis-like conditions associated with infections such as pneumonia and COVID-19.

## ACKNOWLEDGMENTS

This paper is dedicated to the memory of our close colleague, Professor Alexander Laman (Box 1).

### Box 1. Alexander Georgievich Laman (1963–2019)

Born on the 23 April 1963 in Ukraine, Alexander Laman entered Moscow State University in 1981 where he studied in the Department of Bioorganic Chemistry, graduating in 1986. His doctoral studies were at the Russian Academy of Sciences (ROS) Institute of Bioorganic Chemistry in Moscow, where he worked on the structure and functions of human genes under the guidance of Eugene D. Sverdlov. He defended his thesis on the topic of ‘Determination of the nucleotide sequence of a gene fragment encoding the alpha subunit of human Na/K-ATPase’ under the direction of Vladimir A. Nesmeyanov. As a postdoctoral scientist he moved to the branch of the Institute of Bioorganic Chemistry in Pushchino, Moscow Region, where he worked on the molecular basis of the innate immune system. Using phage display, he discovered a peptide mimetic of MDP/GMDP, and using this mimetic uncovered the role of YB-1 in immune signaling by MDP/GMDP. Prof. Laman died on the 7 March 2019 following a short illness at the tragically early age of 56 years.

Box 1 to be placed after the references

